# Genomic analysis of *Cryaa*-R49C and *Cryab*-R120G knockin mutant mice: 10-year follow-up

**DOI:** 10.1101/2020.10.16.343186

**Authors:** Fred Kolling, Carol Ringelberg, Mia Wallace, Usha Andley

**Affiliations:** DartMouse, Mouse Speed Congenic Core Facility, Geisel School of Medicine at Dartmouth, Lebanon, NH 03756, USA; Departments of Pediatrics Washington University School of Medicine, St. Louis, MO 63110, USA; Department of Ophthalmology and Visual Sciences, Washington University School of Medicine, St. Louis, MO 63110, USA

**Author notes:** Correspondence: Usha P. Andley, Ph.D. Department of Ophthalmology and Visual Sciences, 660 South Euclid Avenue, Campus Box 8096, Washington University School of Medicine, St. Louis MO 63110 USA. Co-author e-mails: Fred Kolling IV; Carol Ringelberg; Mia Wallace.

**Keywords:** Mouse genome, SNP markers, cryaa, cryab, mutation, knockin mice

## Abstract

**Objective:** Two experimental samples from a mouse line containing Cryaa or Cryab modifications on a predominantly C57Bl/6 background using 129Sv mouse strain embryonic stem cells were investigated. The objective was to reexamine the precise genetic background of the mice 10 years after they were converted to the C57Bl/6 background.

**Results:** The genetic backgrounds of the mice were assessed at the DartMouse™ Speed Congenic Core Facility at the Geisel School of Medicine at Dartmouth. DartMouse used the Illumina, Inc. Infinium Genotyping Assay to interrogate a custom panel of 5307 SNPs that were spread throughout the genome. The raw SNP data were analyzed using the DartMouse SNaP-Map™ and Map-Synth™ software, which allowed for genetic background identification at each SNP location for every mouse. As part of the analysis, 323 SNPs were eliminated from the data prior to generating chromosome maps due to internal quality control protocols. Of the remaining 4984 SNPs, 44.56% were uninformative (not polymorphic between the two relevant genetic backgrounds) and approximately 0.91% gave uninterpretable data. The remaining 54.53% of the returned SNPs were well distributed throughout the genome. The genetic backgrounds were determined to be 98-99% C57Bl/6J, which was the desired background.

## Introduction

Human cataracts are associated with crystallin gene mutations. Mouse models have been extensively used to investigate cataract etiology, mechanism, and phenotype. Many gene targeting models employ embryonic stem cells that are derived from the 129Sv mouse strain, which carries a mutation in the phakinin gene (CP49). CP49 and its interaction partner, filensin, produce the lens fiber cell-specific intermediate filament known as the beaded filament (1,2). Importantly, the mutation in CP49 that is carried by the 129Sv strain is associated with beaded filaments (3, 4). Experiments with CP49-deleted, knockout mice (5) demonstrated that an absence of beaded filaments leads to reduced lens optical clarity (2). Additionally, the cytoskeletal elements found in the lens including actin, vimentin, and intermediate filaments are known to interact with α-crystallin, which is a molecular chaperone that is abundantly expressed in the lens (6). The aggregation of cytoskeletal proteins such as actin, vimentin, GFAP, and tubulin is inhibited by α-crystallin (6).

To study the effects of *Cryaa* and *Cryab* mutations on cataract formation, we generated mice using 129Sv mouse embryonic stem cells (7, 8). The mice were then initially bred with C57Bl/6J mice for 3 generations to generate a colony with a mixed C57Bl/6J/129Sv background. Subsequently, we used speed congenics to convert the *Cryaa* and *Cryab* knockin mice to the C57Bl/6J background. Analyses that were performed 10 years ago found >95% C57 background in these mice. Here, we characterized the same mice after numerous generations of breeding with C57Bl/6J strain mice. Chromosomal analysis of the genomes from two *Cryaa* and *Cryab* heterozygous knockin mice from our present colony are described. Their genomes were characterized and a high percentage of C57 markers persisted (98-99%).

## Methods

Mice were backcrossed for 5 generations using a marker assisted selection (i.e. “speed congenic”) approach. The mouse genomes were assessed at the DartMouse™ Speed Congenic Core Facility at the Geisel School of Medicine at Dartmouth. DartMouse used the Illumina, Inc. (San Diego, CA) Infinium Genotyping Assay to interrogate a custom panel of 5307 SNPs that were spread throughout the genome. The raw SNP data were analyzed using the DartMouse SNaP-Map™ and Map-Synth™ software, which determined the genetic background at each SNP location for every mouse. The genetic background at the final back-cross generation was determined.

Starting in 2008, the *Cryaa*-R49C and *Cryab*-R120G mouse lines have been propagated at the Mouse Genetics Core in the Department of Pediatrics at Washington University School of Medicine. Mice were either mated with siblings of the same genotype, or with C57Bl/6J WT mice. In 2015, the *Cryaa*-R49C and *Cryab*-R120G mice were re-derived from frozen sperm because we suspected a viral infection in the mouse colonies at the mouse housing facility (based on PCR assays on sentinel mice at the facility). Mice were euthanized by CO_2_ inhalation.

Raw data were used to create a detailed chromosome map for each mouse. The chromosome maps were then used to identify the genetic background of each SNP loci throughout the genome using SNaP-Map software. Map-Synth software was used to graphically display the percentage of the mouse genomes that corresponded to each genetic background. From these graphs and maps, the genetic purity of the speed congenic projects that generated our mouse line was verified (background checks).

All raw SNP genotyping data are presented in an excel file in Supplementary materials (Table S1).

## Results

In the current study, we analyzed two mice. Mouse 1336 had one WT *Cryaa* allele and one *Cryaa*-R49C mutant allele. The second mouse (1396) had one WT *Cryab* allele and one *Cryab*-R120G mutant allele. We previously reported the effects of these mutations on cataract formation (7–11).

For analysis purposes, we assumed that the genetic background of our experimental samples was a mix of C57Bl/6 and 129Sv with no other contaminating genetic backgrounds. In addition, 323 SNPs were eliminated from our data prior to generating chromosome maps due to internal quality control measures. Of the remaining 4984 SNPs, 44.56% were uninformative (i.e. not polymorphic between the two relevant genetic backgrounds) and approximately 0.91% yielded uninterpretable data. The remaining 54.53% of SNPs returned useful data.

Figure 1 displays the chromosome maps of C57Bl/6J, 129Sv, *Cryaa*-R49C-het knockin, and *Cryab*-R120G-het knockin mice. Our analysis showed that these mice had 99.6% and 98.7% C57Bl/6 genomes (Figure 2). Interestingly, the segments of the genomes that showed the highest 129Sv background were located on chromosomes where the mutation was created. For example, in the case of the *Cryaa*-R49C-het mouse 1336, the areas indicated by the purple bands occurred on chromosome 17, which is the mouse chromosome where the *Cryaa* gene is located. Similarly, in the *Cryab*-R120G-het mouse 1396, the purple bands, which indicate the 129Sv and C57Bl backgrounds, occurred on chromosome 9, where the *Cryab* gene is located in the mouse genome.

**Figure 1.**
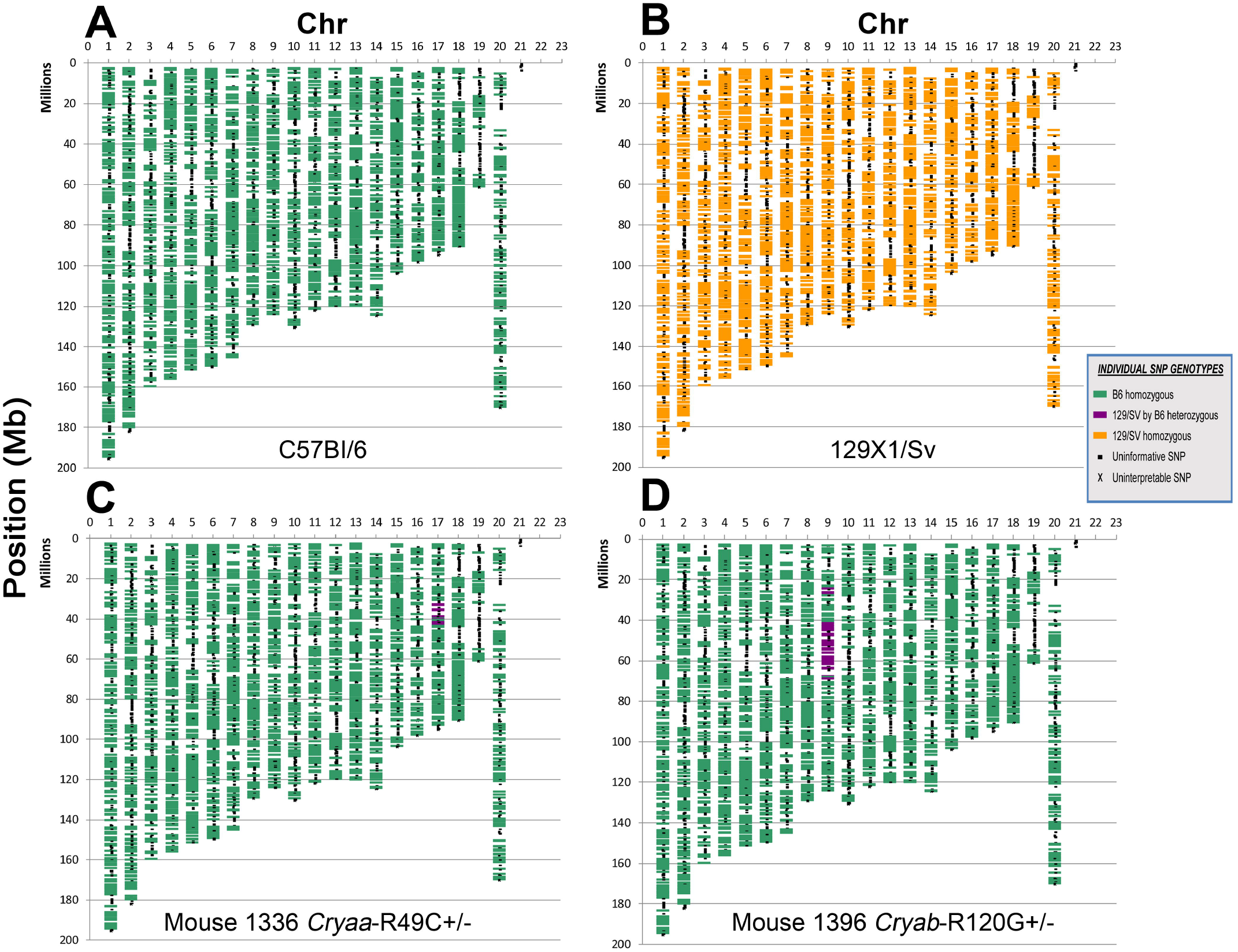
Chromosome maps displaying areas of C57Bl/6 and 129Sv heterozygosity and homozygosity in *Cryaa*-R49C and *Cryab*-R120G mice. Chromosome numbers are shown along the top of each map. Chromosome 20 and 21 are the sex chromosomes (X and Y respectively). Each of the SNP markers in their physical location within the genome of the mouse is shown. These SNP markers are color coded to visually depict the genetic origin as either the “Donor” strain or the “Recipient” strain. *Green*, C57Bl/6: *Purple*, Heterozygous (C57Bl/6 x 129Sv): *Orange:* 129Sv. The knockin modification can be clearly identified on the proximal ends of chromosomes 17 and 9 for the *Cryaa*-R49C and *Cryab*-R120G mice, respectively.

This type of analysis demonstrates the power of the SNaP-Map software for assigning each marker to the proper genetic background. The current analysis duplicates a similar analysis that was performed in 2010 by DartMouse on earlier generations of the same mouse lines. In the 2010 analysis, after backcrossing and speed congenics, the C57Bl/6 background was 97% in *Cryaa*-R49C het mice and 95.8% in *Cryab*-R120G-het mice. The repeated analysis in 2019, after multiple breeding cycles and re-derivation from frozen sperm, indicated that the C57Bl/6 background percentage had increased by 2-3%. The background was 99.6% and 98.7% C57Bl/6 for Cryaa-R49C-het and Cryab-R120G-het mice, respectively (Figure 2A).

**Figure 2.**
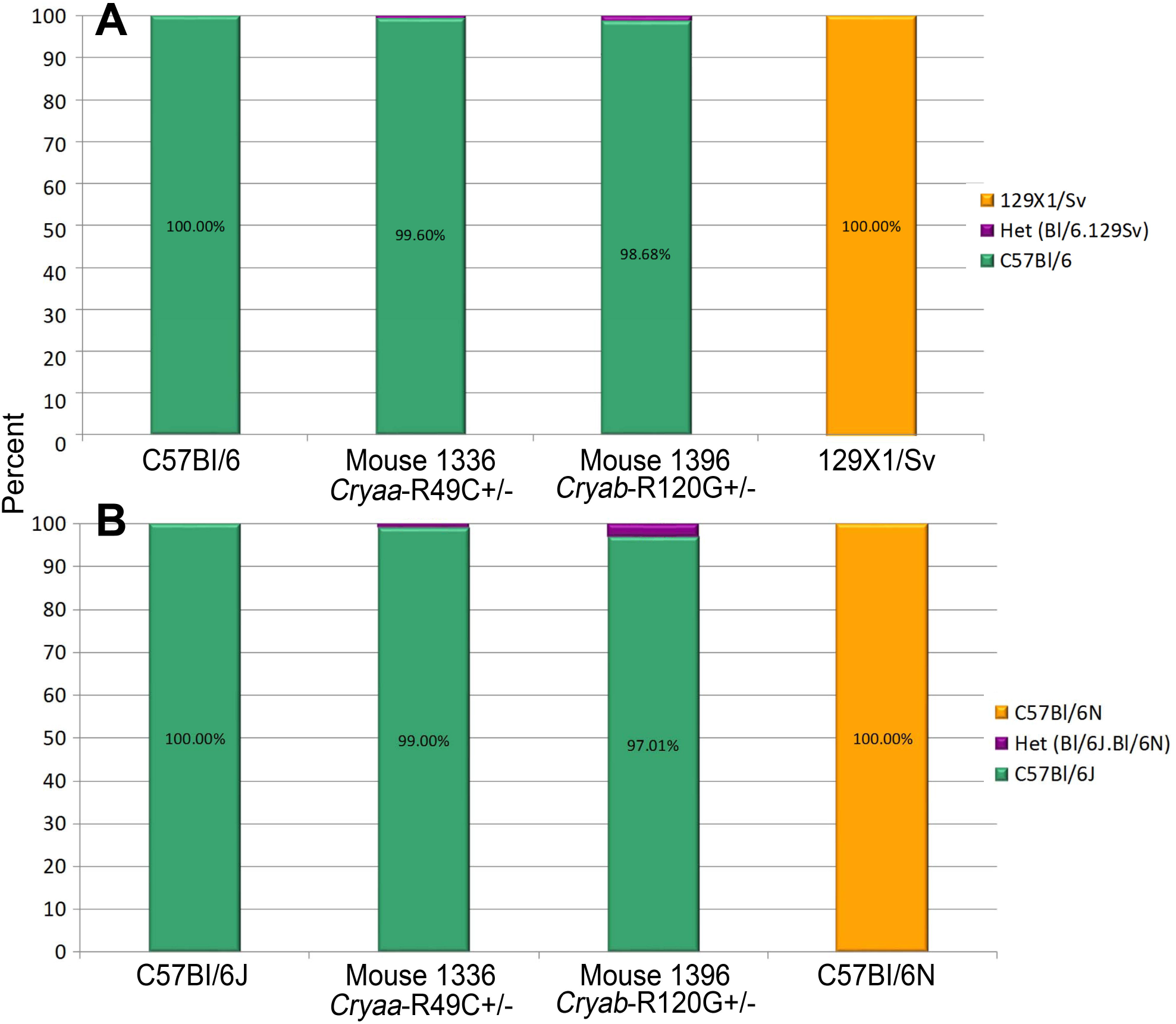
Graphs displaying genetic material percentages. *(A)* C57Bl/6 or 129Sv in *Cryaa*-R49C mouse 1336 and *Cryab*-R120G mouse 1396. *(B)* C57Bl/6J or C57Bl/6N in *Cryaa*-R49C mouse 1336 and *Cryab*-R120G mouse 1396. (Part of the graph in *(A)* was published in Invest Ophthalmol Vis Sci. 2019: 60: 3320–3331. Copyright Association for Research in Vision and Ophthalmology. https://doi.org/10.1167/iovs.18-25647).

Further analysis of the same data showed that the *Cryaa*-R49C-het knockin and *Cryab*-R120G-het knockin mice had 99.0% and 97.01% C57Bl/6J backgrounds (Figure 2B). A comparison of the C57Bl/6J and C57Bl/6N chromosome maps is displayed in Figure 3. The full SNP panel contained 201 distinct markers, which distinguished the C57Bl/6 “J” from the C57Bl/6 “N” sub-strains. We analyzed these SNP markers to determine the J and N strain percentages in the Cryaa and Cryab knockin mice. Chromosome maps that include these SNPs were analyzed to distinguish between the B6J and B6N substrains (Figure 3).

**Figure 3.**
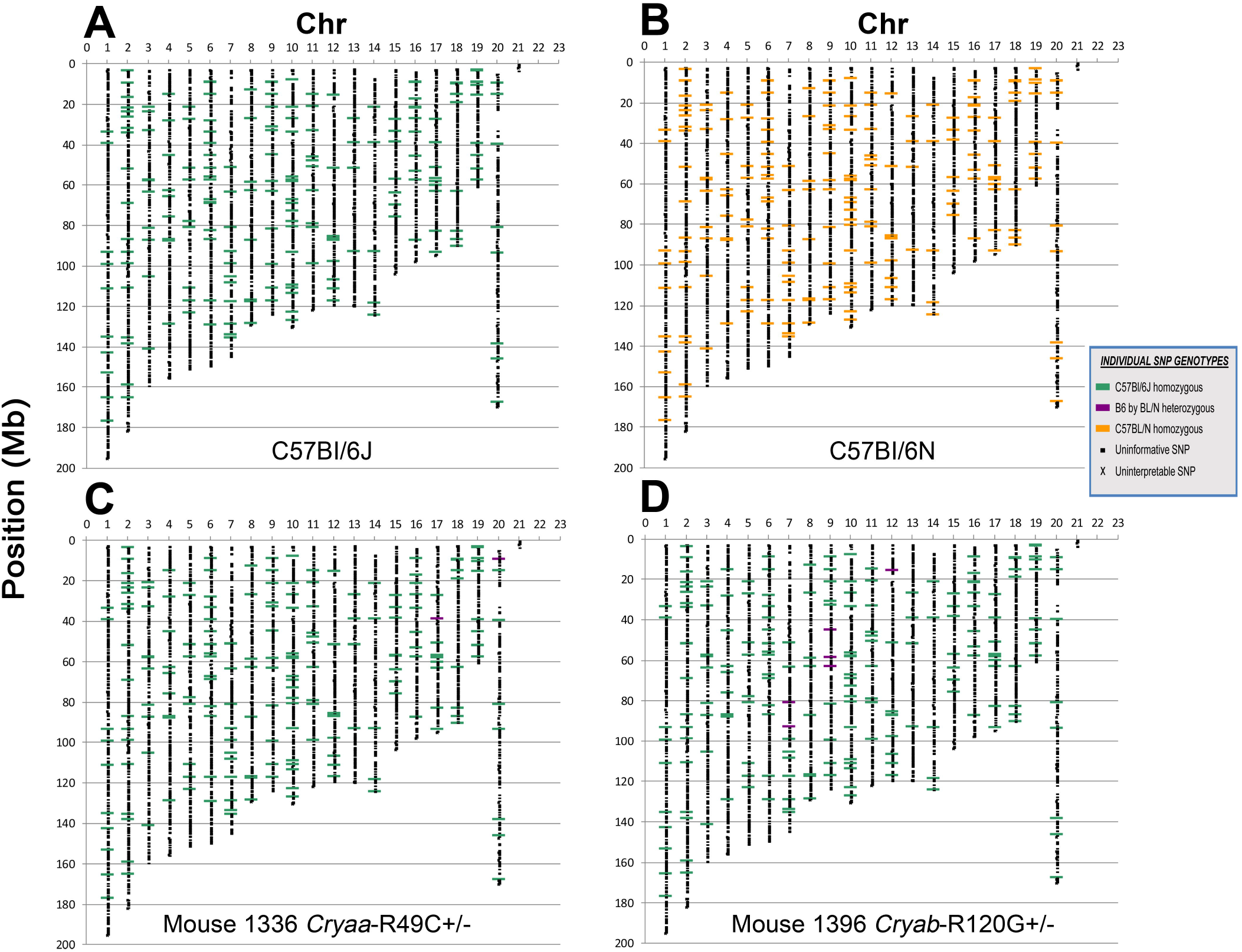
Maps displaying areas of C57/Bl6J and C57Bl/6N heterozygosity and homozygosity in *Cryaa*-R49C and *Cryab*-R120G mice. Each of the SNP markers in their physical location within the mouse genome is shown. These SNP markers are color coded to visually depict the genetic origin as either the C57Bl/6J or C57Bl/6N strain. *Green*, C57Bl/6J homozygous; *Purple*, Heterozygous (C57Bl/6 x C57Bl/6N): *Orange:* C57Bl/6N homozygous.

Of the 5307 loci on our SNP panel, 83 fell outside of the DartMouse 95% quality control confidence interval and were not referenced in the analyses. Additionally, as displayed in Table S2 (Supplementary materials), a small percentage of the SNPs that were interrogated returned uninterpretable data. Historically, approximately 1% of the total panel of SNPs that are investigated using DartMouse protocols return non-useable data. These were located throughout the genome and are shown in Figures 1 and 3.

## Discussion

The 129Sv mouse strain carries a mutation in the gene for CP49 (phakinin) and also lacks CP49, which is an intermediate filament protein that is specific to lens fiber cells. Consequently, 129Sv mice are almost completely deficient in CP49 protein and its interaction partner, filensin, which results in the absence of the beaded filament in the lens. The 129Sv mouse strain exhibits a slow and progressive loss of optical clarity. In addition, the lens cytoskeleton interacts with αB-crystallin, and therefore, the absence of the beaded filament can confound interpretations of αB-crystallin mutation effects. Thus, we sought to convert these mice to the C57Bl/6J background, which possesses a normal CP49 gene level (4).

Many knockout and knockin mouse models utilize embryonic stem cells from the 129Sv strain. This can confound the interpretation of cataracts that occur as a result of other gene mutations and is especially complicated when exploring genes, including crystallin, that are known to interact with lens beaded filaments.

Here, we sought to establish the C57Bl/6J background in mice that had mutations in the human cataract-linked Cryaa and Cryab genes. These mice were backcrossed with C57BL/6J mice by speed congenics. Lens fiber cells that lack CP49 are more pliable than stiffer fiber cells derived from WT mice. The cells that lack CP49 (CP49 mutant) also have a higher axial to equatorial diameter ratio (3). Notably, 129Sv embryonic stem cells are often used in gene targeting studies even though the presence of a CP49 mutation confounds the cataract phenotype. Although the lens fiber cells in the 129Sv strain mice initially appear to differentiate and elongate normally, the 129Sv phenotype develops slowly, and a progressive loss of lens optical clarity is detectable by electron microscopy.

The absence of CP49, the near absence of filensin, and the resultant absence of the beaded filament may have downstream effects on any proteins or structures that are binding partners with these proteins. Thus, the true ramifications of the CP49 mutation in past gene-targeting studies may be difficult to assess. To eliminate the possibility that the CP49 mutation is affecting the results, it may be necessary to backcross the target gene to a wild-type background.

The speed congenics methods that we describe, combined with SNP and chromosome analyses, can be applied to other mouse models. An *Rd8* mutation has been found in the C57Bl/6N mouse strain and produces retinal phenotypes (12). The speed congenics method used in this study could be useful to convert those mice to a more suitable genetic background. Using genomic analysis of the Cryaa and Cryab knockin mouse chromosomes, we demonstrated that these mice have C57 characteristics instead of 129Sv strain traits. The genomes were 97 and 99% C57Bl/6J. Thus, cataracts observed in these knockin mice are not confounded by the presence of the phakinin mutation and are exclusively due to Cryaa or Cryab mutations.

The 129Sv mouse strain is widely used as a source of embryonic stem cells for generating knockin and knockout mouse models for cataract lens studies. Starting in 2007, we backcrossed knockin mice that had *Cryaa* or *Cryab* mutations with C57Bl/6J mice for 3 generations. Subsequently, we used speed congenics in 2009-2010 to generate mice that had a higher purity C57Bl/6J background. Here, we report the same analysis that was completed in 2010, but are updating the analysis in 2019 to include mice that were sib-mated for many generations over the years. The C57Bl/6J background mouse models still produced cataracts, which suggests that the cataracts are unlikely to be caused by the CP49 mutation.

## Limitations

We assumed that the genetic background of the experimental samples was a mix of C57Bl/6 and 129Sv, with no other contaminating backgrounds.

## Supporting information

Animal procedures checklist

## List of Abbreviations

SNP: single-nucleotide polymorphism;
WT: Wild-type

## Declarations

### Ethics approval and consent to participate

All animal experiments were approved by the Washington University in St. Louis IACUC Committee (Protocol number 20170212). No human studies were conducted in this research.

### Consent for publication

Each author has agreed to publication of this manuscript.

### Availability of data and materials

All data generated or analyzed during this study are included in the published article and supplementary information files.

### Authors’ contributions

**FK:** Data acquisition, data analysis, data interpretation, report writing: **CR:** Data acquisition, data analysis, data interpretation, report writing: **MW:** Conception and design of the work, manuscript writing: **UPA:** Manuscript writing, funding procurement.

All authors have approved the submitted version of the manuscript and have agreed to be both personally accountable for their own contributions and to ensure that questions related to the accuracy or integrity of any part of the work, even ones in which the author was not personally involved, are appropriately investigated and resolved and that the resolution is documented in the literature.

## Acknowledgements

Not applicable.

## Supplementary Material

**Table S1:** Binary raw data interrogated from the analysis of *Cryaa*-R49C and *Cryab*-R120G mice.

**Table S2:** Summary of SNP analysis of *Cryaa*-R49C and *Cryab*-R120G mice.

