## Supplemental Table 1 for "Genomic analysis of *Cryaa*-R49C and *Cryab*-R120G knockin mutant mice: 10-year follow-up"

**Supplementary Material. Table S1.**

Table S1: Summary of SNP Analysis

Total number of samples analyzed

2

Analysis Type

Genetic Background Check

Analysis Platform

5307 SNP Illumina Infinium BeadChip

DartMouse SNaP

-

Map Software

SNP Information

2221 Uninformative (44.56%)

2718 Informative (54.53%)

SNP Quality C

ontrol

0.91% (~45 total) of SNPs were

uninterpretable;

0 non

-

control

-

like SNPs

Inbred Strains

C57Bl/6 and 129Sv Inbred Strains

Genetic Modification

Modificatio

n of the Cryaa or Cryab

genes
