## Supplementary material for "Genomic analysis of *Cryaa*-R49C and *Cryab*-R120G knockin mutant mice: 10-year follow-up": Animal procedures checklist

### The ARRIVE guidelines 2.0: author checklist

#### The ARRIVE Essential 10

These items are the basic minimum to include in a manuscript. Without this information, readers and reviewers cannot assess the reliability of the findings.

| Item | | Recommendation | Section/line number, or reason for not reporting |
|---|---|---|---|
| **Study design** | 1 | For each experiment, provide brief details of study design including:<br>a. The groups being compared, including control groups. If no control group has been used, the rationale should be stated.<br>b. The experimental unit (e.g. a single animal, litter, or cage of animals). | WT, Cryaa-R49C, Cryab-R120G, C57Bl/6J, 129Sv<br>Two animals |
| **Sample size** | 2 | a. Specify the exact number of experimental units allocated to each group, and the total number in each experiment. Also indicate the total number of animals used.<br>b. Explain how the sample size was decided. Provide details of any *a priori* sample size calculation, if done. | One<br>Not applicable |
| **Inclusion and exclusion criteria** | 3 | a. Describe any criteria used for including and excluding animals (or experimental units) during the experiment, and data points during the analysis. Specify if these criteria were established *a priori*. If no criteria were set, state this explicitly.<br>b. For each experimental group, report any animals, experimental units or data points not included in the analysis and explain why. If there were no exclusions, state so.<br>c. For each analysis, report the exact value of *n* in each experimental group. | Heterozygous mice<br>Not applicable<br>1 |
| **Randomisation** | 4 | a. State whether randomisation was used to allocate experimental units to control and treatment groups. If done, provide the method used to generate the randomisation sequence.<br>b. Describe the strategy used to minimise potential confounders such as the order of treatments and measurements, or animal/cage location. If confounders were not controlled, state this explicitly. | No<br>Not applicable |
| **Blinding** | 5 | Describe who was aware of the group allocation at the different stages of the experiment (during the allocation, the conduct of the experiment, the outcome assessment, and the data analysis). | Not applicable |
| **Outcome measures** | 6 | a. Clearly define all outcome measures assessed (e.g. cell death, molecular markers, or behavioural changes).<br>b. For hypothesis-testing studies, specify the primary outcome measure, i.e. the outcome measure that was used to determine the sample size. | Not applicable<br>Not applicable |
| **Statistical methods** | 7 | a. Provide details of the statistical methods used for each analysis, including software used.<br>b. Describe any methods used to assess whether the data met the assumptions of the statistical approach, and what was done if the assumptions were not met. | Not applicable<br>Not applicable |
| **Experimental animals** | 8 | a. Provide species-appropriate details of the animals used, including species, strain and substrain, sex, age or developmental stage, and, if relevant, weight.<br>b. Provide further relevant information on the provenance of animals, health/immune status, genetic modification status, genotype, and any previous procedures. | These studies used mice of the strain<br>C57BL/6J background, [illegible] |
| **Experimental procedures** | 9 | For each experimental group, including controls, describe the procedures in enough detail to allow others to replicate them, including:<br>a. What was done, how it was done and what was used.<br>b. When and how often.<br>c. Where (including detail of any acclimatisation periods).<br>d. Why (provide rationale for procedures). | Animals were euthanized prior to experiments. [illegible]<br>Chromosomes were<br>DartMouse<br>To otain genomic |
| **Results** | 10 | For each experiment conducted, including independent replications, report:<br>a. Summary/descriptive statistics for each experimental group, with a measure of variability where applicable (e.g. mean and SD, or median and range).<br>b. If applicable, the effect size with a confidence interval. | Not applicable<br>Not applicable |

### The Recommended Set

These items complement the Essential 10 and add important context to the study. Reporting the items in both sets represents best practice.

| Item | | Recommendation | Section/line number, or reason for not reporting |
|---|---|---|---|
| **Abstract** | 11 | Provide an accurate summary of the research objectives, animal species, strain and sex, key methods, principal findings, and study conclusions. | Two experimental samp |
| **Background** | 12 | a. Include sufficient scientific background to understand the rationale and context for the study, and explain the experimental approach. | Here, we sought to establish the C57Bl/6J [illegible] |
| | | b. Explain how the animal species and model used address the scientific objectives and, where appropriate, the relevance to human biology. | The mice are being used as model [illegible] |
| **Objectives** | 13 | Clearly describe the research question, research objectives and, where appropriate, specific hypotheses being tested. | The objective was to re |
| **Ethical statement** | 14 | Provide the name of the ethical review committee or equivalent that has approved the use of animals in this study, and any relevant licence or protocol numbers (if applicable). If ethical approval was not sought or granted, provide a justification. | IACUC, Washington University in St. Louis. Protocol number 20170212 |
| **Housing and husbandry** | 15 | Provide details of housing and husbandry conditions, including any environmental enrichment. | Division of Comparative |
| **Animal care and monitoring** | 16 | a. Describe any interventions or steps taken in the experimental protocols to reduce pain, suffering and distress. | Animals were euthanized prior to [illegible] |
| | | b. Report any expected or unexpected adverse events. | None |
| | | c. Describe the humane endpoints established for the study, the signs that were monitored and the frequency of monitoring. If the study did not have humane endpoints, state this. | N/A |
| **Interpretation/ scientific implications** | 17 | a. Interpret the results, taking into account the study objectives and hypotheses, current theory and other relevant studies in the literature. | Using genomic analysis of the Cryaa [illegible] |
| | | b. Comment on the study limitations including potential sources of bias, limitations of the animal model, and imprecision associated with the results. | We assumed that the genetic background of [illegible] |
| **Generalisability/ translation** | 18 | Comment on whether, and how, the findings of this study are likely to generalise to other species or experimental conditions, including any relevance to human biology (where appropriate). | The C57Bl/6J background mouse models still produced [illegible] |
| **Protocol registration** | 19 | Provide a statement indicating whether a protocol (including the research question, key design features, and analysis plan) was prepared before the study, and if and where this protocol was registered. | Mouse Genetics Core, Washington University |
| **Data access** | 20 | Provide a statement describing if and where study data are available. | Manuscript is submitted |
| **Declaration of interests** | 21 | a. Declare any potential conflicts of interest, including financial and non-financial. If none exist, this should be stated. | None |
| | | b. List all funding sources (including grant identifier) and the role of the funder(s) in the design, analysis and reporting of the study. | This work was supported by the [illegible] |
